# Sparcle: assigning transcripts to cells in multiplexed images

**DOI:** 10.1101/2021.02.13.431099

**Authors:** Sandhya Prabhakaran, Tal Nawy, Dana Pe’er’

## Abstract

**Background:** Imaging-based spatial transcriptomics has the power to reveal patterns of single-cell gene expression by detecting mRNA transcripts as individually resolved spots in multiplexed images. However, molecular quantification has been severely limited by the computational challenges of segmenting poorly outlined, overlapping cells, and of overcoming technical noise; the majority of transcripts are routinely discarded because they fall outside the segmentation boundaries. This lost information leads to less accurate gene count matrices and weakens downstream analyses, such as cell type or gene program identification.

**Results:** Here, we present Sparcle, a probabilistic model that reassigns transcripts to cells based on gene covariation patterns and incorporates spatial features such as distance to nucleus. We demonstrate its utility on both multiplexed error-robust fluorescence in situ hybridization (MERFISH) and single-molecule FISH (smFISH) data.

**Conclusions:** Sparcle improves transcript assignment, providing more realistic per-cell quantification of each gene, better delineation of cell boundaries, and improved cluster assignments. Critically, our approach does not require an accurate segmentation and is agnostic to technological platform.

## Introduction

Imaging-based spatial transcriptomics comprises a set of ground-breaking technologies that scale up the number of genes that can be quantified within tissues (1–6). The emergence of these approaches has made it possible to resolve cell type heterogeneity and to gain a holistic understanding of gene expression within spatial context (7–9). However, computational tools to interpret this data are severely lacking. A fundamental challenge is the construction of a biologically accurate cell-by-gene count matrix, which requires cell segmentation as a first step (10). Differences in cell size and morphology, occlusion, overlapping cells, de-nucleated cells and technical noise from the imaging instruments make simultaneous automated segmentation of multiple cell types extremely difficult.

The boundaries of nuclei are easier to resolve than cell outlines. Current automated cell segmentation methods identify nuclei using data from a nuclear marker channel such as DAPI, then radially dilate these borders by a few pixels to approximate cell boundaries. This tends to leave a substantial fraction of ‘dangling’ transcripts outside of defined cell boundaries (**Fig. 1**). Dangling transcripts are potentially assignable to multiple cells, especially as distance from the nucleus increases, and are thus discarded prior to generating the count matrix. Moreover, 2D images only reveal a slice of the cell volume. The transcripts present in peri-nuclear sections therefore represent a very sparse sampling of the cellular transcriptome, which underpowers downstream analyses of cell types and gene programs.

**Figure 1:**
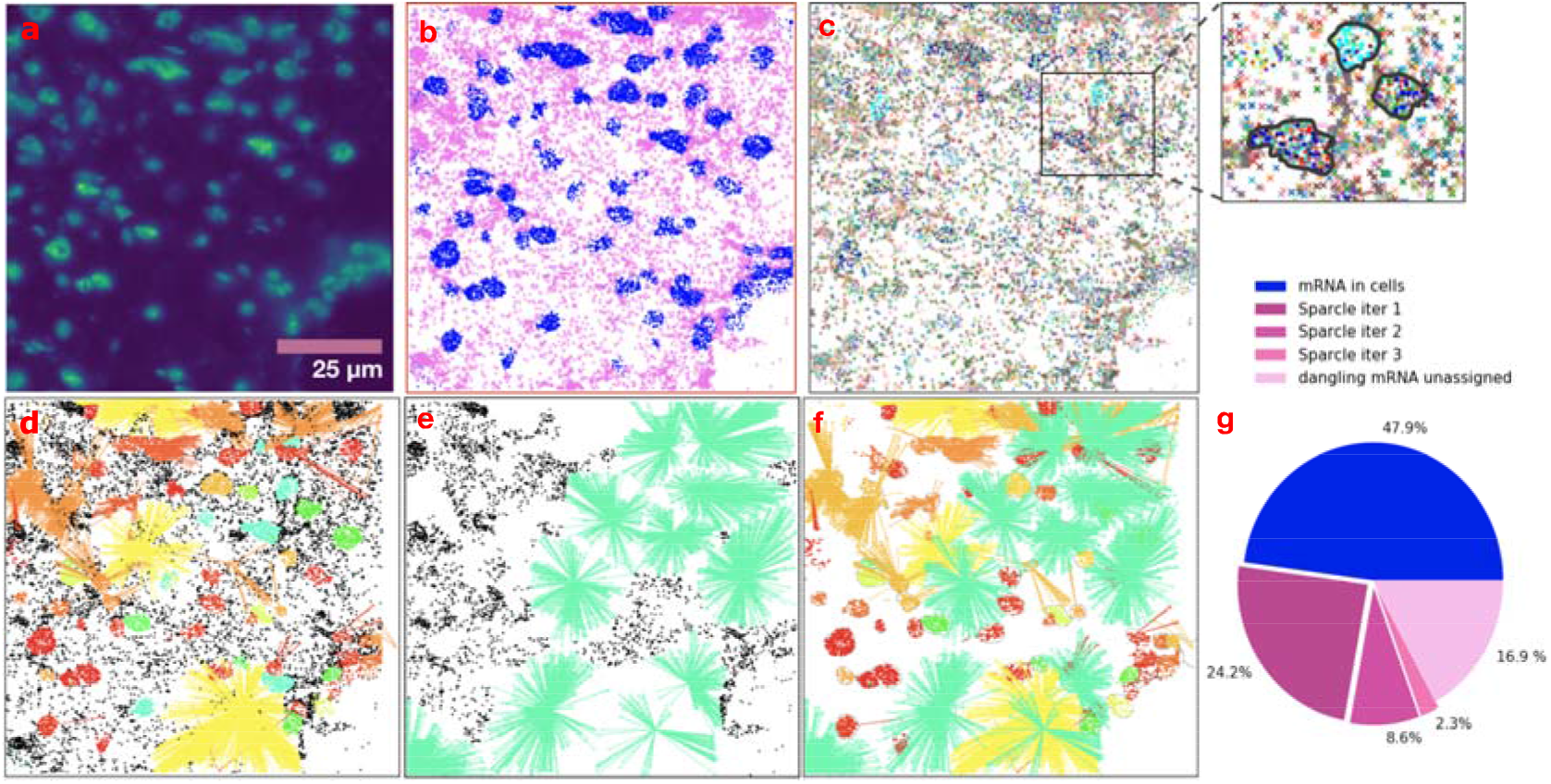
Sparcle captures transcriptomic diversity that is present in spatial transcriptomic images but is discarded by typical analysis. **a–f**. A single 2D exemplar field of view (FOV) of the mouse hypothalamic posterior optic region imaged by multiplexed error-robust fluorescence in situ hybridization (MERFISH) [9]. **a**. MERFISH image showing individual transcripts as points of green fluorescence. **b**. Conservative segmentation captures n = 12,000 transcripts (blue dots; ~48% of mRNAs) (Moffitt et al., *Science* 2018), leaving ~14,000 ‘dangling’ transcripts (pink crosses). Current methods fail to accurately segment neuronal cell morphologies in particular, which often contain transcripts far from the cell body. **c**. Spatial diversity of transcripts within (dots) and outside (crosses) the cell segmentation. Each mRNA is colored according to corresponding gene (n = 140 genes assayed). Inset, magnification highlighting the diversity of dangling transcripts; solid lines indicate segmentation boundaries**. d–f**. For the FoV shown in (**a**), Sparcle captures 27%, 10.3% and 1.2% of total mRNAs in iteration 1 (**d**), iteration 2 (**e**) and iteration 3 (**f**), respectively. (Panel (**f**) also overlays all the mRNA assignments from previous iterations). **f**. Overall dangling mRNA assignments across the three iterations, with unassigned dangling mRNAs removed. Dangling mRNAs assigned to one cluster are connected to the cell centroid by lines of the cluster color; black dots are unassigned dangling mRNAs. **g**. Fraction of all mRNAs recovered across 400 MERFISH FOVs by segmentation and Sparcle assignment.

To address these issues, we introduce Sparcle (spatial reassignment of spots to cells via maximum likelihood estimation), a comprehensive probabilistic model that assigns dangling transcripts on the basis of gene covariance, as well as the transcripts’ distance between neighboring cells and adjacent transcripts. These patterns include neighboring cells (defined by Euclidean distances between cell centroids), and neighboring transcripts for each cell. Sparcle generalizes to any smFISH imaging methodology, and we demonstrate its utility on both multiplexed error-robust fluorescence in situ hybridization (MERFISH) (9) and single-molecule FISH (smFISH) (5, 8) data. Sparcle is one of the first methods to capture dangling transcripts; we show that it recovers more than 50% of these mRNAs, and validate its performance using scRNA-seq data.

## Results

### Sparcle model

Sparcle is a probabilistic model that iteratively assigns dangling transcripts to neighboring cells using maximum likelihood estimation (Methods). Importantly, our approach does not require a highly accurate segmentation as input.

The large number of transcripts in each cell justifies modelling gene expression as a multivariate Gaussian distribution, based on the central limit theorem assumption. An image can thus be considered as a mixture of cell-specific Gaussian distributions, which can be posed in a Dirichlet process mixture model (DPMM) (11–13) setting to identify heterogeneous cell types. In Sparcle, the user can choose either the DPMM or Phenograph (14) for clustering; neither require that cluster numbers are known a priori. We then compute a multivariate Gaussian distribution representing gene-gene covariation relationships for each cluster in our data by learning the cluster-specific first and second order moments (mean and covariance) of each cluster. For every dangling mRNA, the algorithm constructs a circular weighted ‘mock’ cell with radius *r*, centered at the mRNA, that captures proximal mRNA neighbors (**Fig. S1** and Methods). The contribution of each gene to the mock cell is inversely weighted by its distance from the dangling mRNA. Next, Sparcle generates a maximum likelihood estimate (MLE) of which cluster (often corresponding to cell type) the mock cell most likely represents and assigns the dangling mRNA to the cell with that cluster label nearest to the mock cell, and updates the count matrix. This process of assigning dangling mRNAs continues for a fixed number of iterations by updating and clustering the count matrix, recalculating the cluster-specific moments and revising the cellular assignments to clusters at each iteration. The number of robustly assigned transcripts per iteration exhibits asymptotic behavior, and we have found little additional benefit beyond the default of 3 iterations. With a user-provided cell segmentation and a minimal set of default parameters, Sparcle can robustly assign dangling mRNAs in under 10 min on a high-performance cluster for a typical MERFISH FOV consisting of ~80 cells and ~14,000 dangling mRNAs.

The outputs of the iterative model are a) a more realistic count matrix, based on improved mRNA assignments to cells, b) cluster assignments for each cell and c) improved delineation of cell boundaries.

### Evaluating Sparcle using matching scRNA-seq data

The mammalian brain is characterized by diverse cell types with complicated morphologies and cellular interactions that can greatly benefit from multiplexed imaging and spatial analysis. A MERFISH image of the mouse hypothalamus posterior optic region comprises ~30,000 cells from 70 different neuronal populations and 140 assayed genes (9) (**Fig. 1**). From this image, 400 smaller fields of view (FOVs) were imaged at high magnification (**Fig. 1a**); segmentation using dilated cell nuclei revealed that ~52% of mRNAs, which exhibit substantial spatial heterogeneity, remain unassigned (**Fig. 1b,c**). Using these individual FOVs (**Fig. 1d-f**), Sparcle increases the fraction of total transcripts that are assigned to cells by assigning 24.2, 8.6 and 2.3% out the total transcripts in iterations 1,2 and 3, respectively (**Fig. 1g**), ultimately capturing 68% of the 2.8 million dangling transcripts in this full dataset.

To evaluate the cell populations identified by Sparcle, we used paired scRNA-seq data (31,299 cells) as ground truth, considering a subset of 900 genes containing MERFISH probes and neuronal genes, as utilized in Moffitt et al. [9], to call cell types. We used Phenograph (14) to cluster both the scRNA-seq and MERFISH count matrices before and after applying Sparcle. Post-Sparcle, we observe that the cell type proportions are more concordant with those of the scRNA-seq data (**Fig. 2a**).

**Figure 2;.**
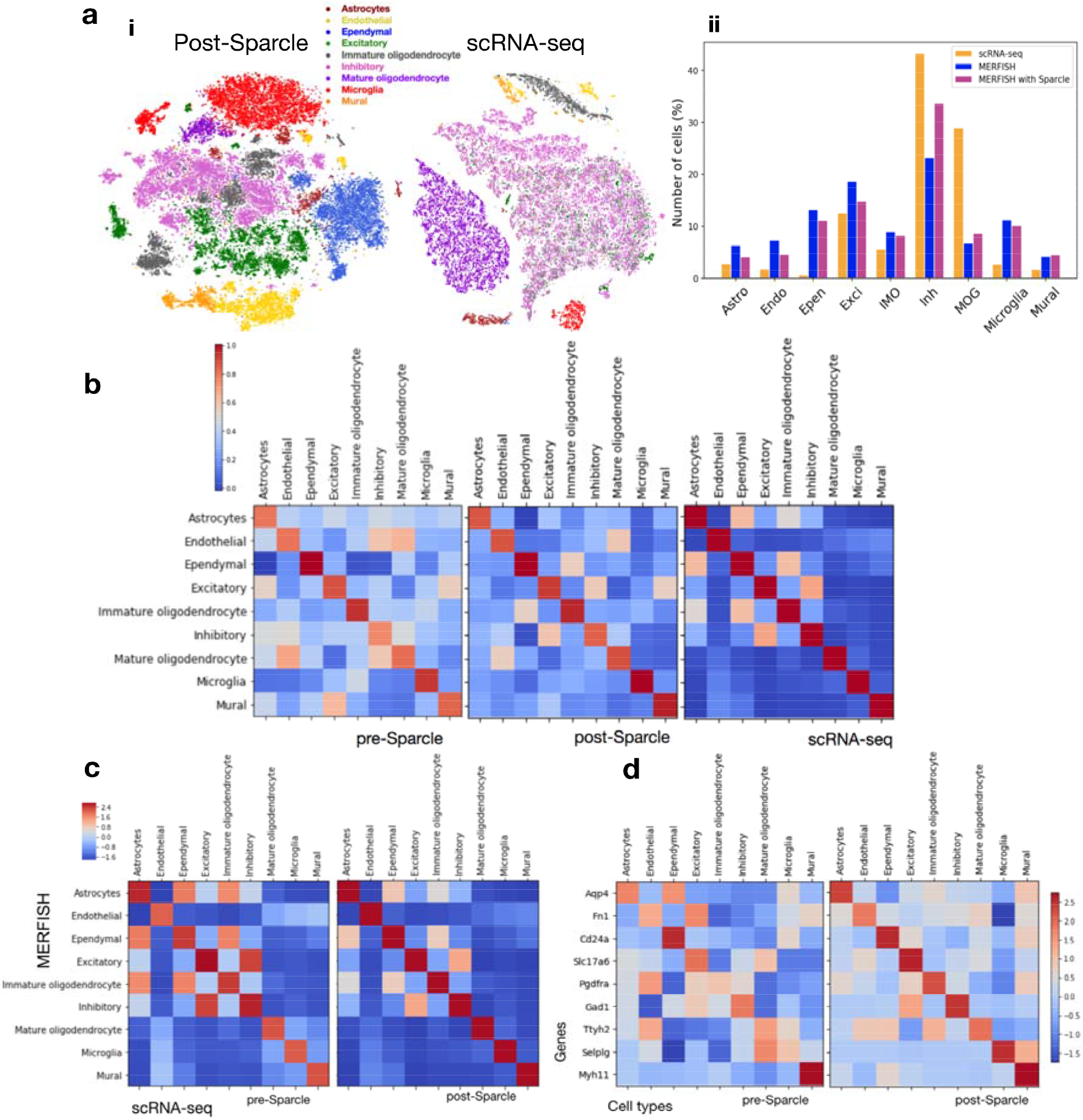
Sparcle improves cell type assignment from MERFISH data. **a**. (i) t-SNE projections of MERFISH image data from the mouse hypothalamus optic region comprising 400 fields of view (FOV) with ~30,000 cells total, 140 genes, with matching scRNA-seq data in Moffitt et al. [9] showing diversity in cell type proportions. (ii) Percentage of cells in the paired scRNA-seq data (orange bars), MERFISH (blue bars) and MERFISH with Sparcle (pink bars). Post-Sparcle cell type assignment proportions are in closer accordance with the scRNA-seq proportions, especially for astrocytes, endothelial cells, mature and immature oligodendrocytes, and excitatory and inhibitory neuronal cell types, demonstrating that Sparcle can correct counts per cell type based on the dangling mRNA assignments. **b**. Cluster-cluster covariance (Gramian) matrices of MERFISH (left), MERFISH with Sparcle (center) and scRNA-seq (right) indicate that Sparcle-treated count matrices are more similar to scRNA-seq (pre-Sparcle vs. scRNA-seq median χ^2^ (2, N = ~61,000) = 0.26, p = 0.31, Frobenius norm = 15.18; post-Sparcle vs. scRNA-seq median χ2 (2, N = ~61,000) = 0.95, p = 0.62, Frobenius norm = 2.34). **c**. Cluster-cluster cross covariance matrices between scRNA-seq and MERFISH clusters pre- and post-Sparcle indicate stronger within-cluster variances (dominant diagonal) after Sparcle, especially for non-neuronal cell types. **d**. Sparcle improves the cell-type specificity of canonical gene assignment for most cell types. z-score-normalized gene expression (rows) is averaged over vectors per cluster representing cell type (columns).

To further assess Sparcle’s performance, we tested the similarity of count matrices derived from MERFISH and scRNA-seq using two different covariance approaches (Methods). The Gramian (cluster-cluster) covariance matrices from scRNA-seq and MERFISH are more similar after Sparcle (p = 0.620, Box’s M test; Frobenius norm = 2.34) than before Sparcle (p = 0.309; Frobenius norm = 15.18; larger p-values and smaller norms indicate more similar matrices), showing that Sparcle assignments improve MERFISH cell type calls relative to ground truth (**Fig. 2b**). The cluster-cluster cross covariance matrix between scRNA-seq and post-Sparcle MERFISH exhibits stronger variances within clusters (dominant diagonal) than between clusters, compared with the pre-Sparcle cross covariance matrix, especially for non-neuronal cell types (**Fig. 2c**). This covariance pattern signals stronger relationships within cell types and weaker relationships between different cell types, supporting the accuracy of Sparcle assignments.

Finally, Sparcle clearly improves the assignment of canonical cell-type-specific genes for most clusters, including excitatory neuronal, endothelial, ependymal and astrocyte cells (**Fig. 2d**). This observation supports the validity of the higher proportions of these cell types that result from Sparcle transcript assignments (**Fig. 2a**).

### Sparcle captures more information in MERFISH-derived cell clusters

We sought to determine whether Sparcle assignments capture more transcriptional variability and lend greater structure to molecular quantification data from imaging datasets. To do this, we examined pairwise Pearson correlation coefficients between cluster expression profiles extracted from the mouse hypothalamus scRNA-seq and MERFISH data, both before and after Sparcle, and used Loewner ordering to compare matrices (see Methods).

We first extracted and examined two broad neuronal cell types. We built a pre-Sparcle MERFISH matrix with 100 randomly sampled excitatory cells stacked over 100 inhibitory cells and a post-Sparcle matrix with the same cells, then computed Pearson correlations with a similarly constructed scRNA-seq matrix of 100 random cells per cell type (**Fig. 3a**). The matrices reveal higher scRNA-seq correlations and greater correlation structure for both cell types after using Sparcle (**Fig. 3a**). Notably, Sparcle assignments substantially improve the recovery of excitatory cells, which only number approximately half of the inhibitory cells. Furthermore, Loewner ordering reveals that the excitatory block in the post-Sparcle correlation matrix captures additional variability (subtracting this block in the post-Sparcle matrix from the pre-Sparcle matrix generates a positive semi-definite matrix, whereas subtraction in the other direction does not).

**Figure 3.**
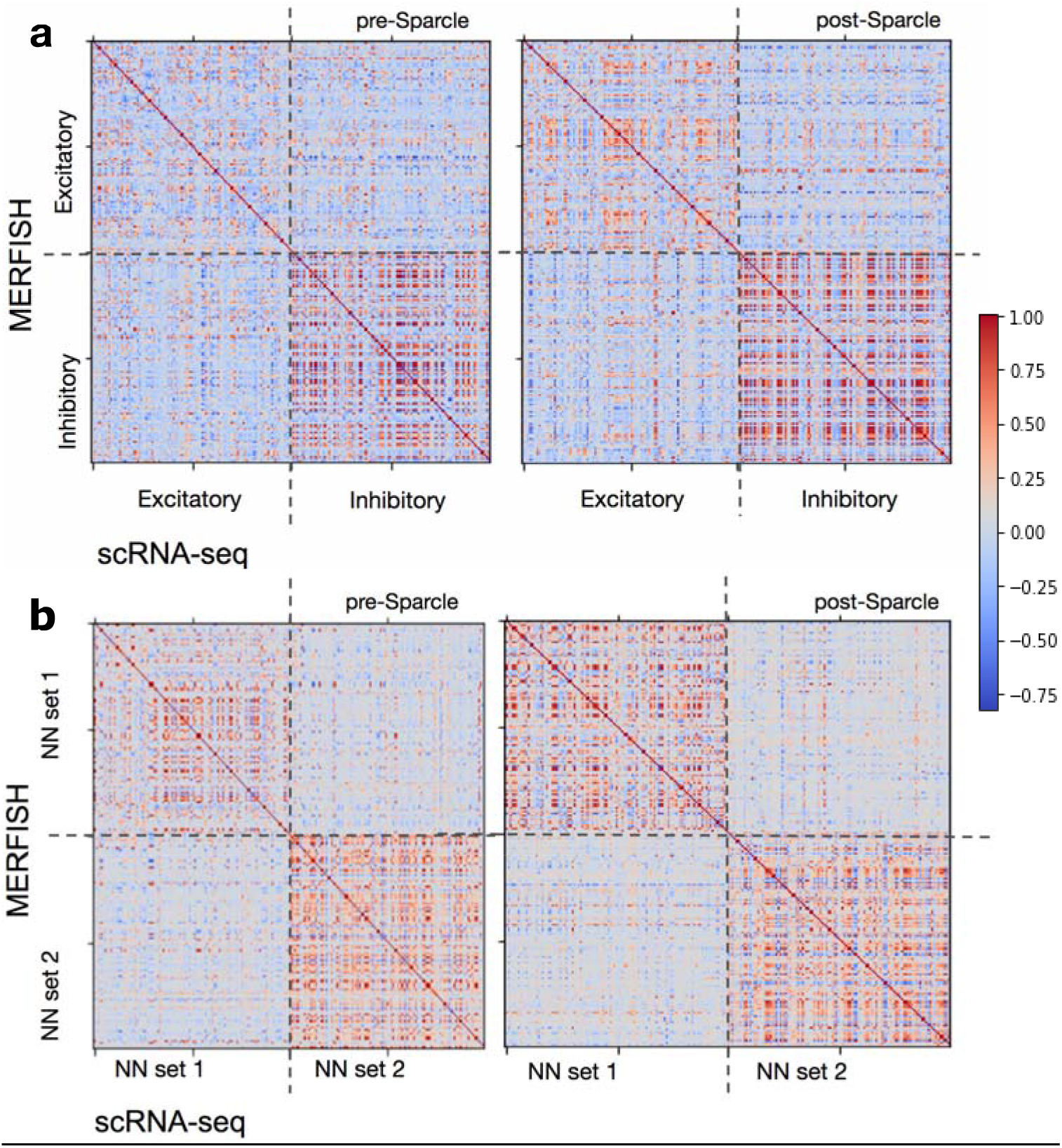
Sparcle refines clusters in MERFISH data. **a.** We constructed a pre-Sparcle MERFISH matrix with 100 randomly sampled excitatory cells stacked over 100 inhibitory cells and a post-Sparcle matrix with the same cells, then computed Pearson correlations with a similarly constructed scRNA-seq matrix of 100 random cells per cell type. Pearson correlation matrices indicate greater covariance across diagonal blocks (defined red zones) between scRNA-seq and post-Sparcle MERFISH data, for both excitatory and inhibitory neuronal cell types. **b.** Similar analysis reveals the same trends for astrocytes, endothelial and ependymal cells (NN set 1) and immature and mature oligodendrocytes, microglia and mural cells (NN set 2).

For non-neuronal clusters, we placed astrocytes, endothelial and ependymal cells in one set (NN set 1) and immature and mature oligodendrocytes, microglia and mural cells in another (NN set 2), and performed Pearson correlation and Loewner ordering (**Fig. 3b**). Both cell sets show clearly higher correlations with scRNA-seq, and the Pearson correlation block diagonals in the post-Sparcle matrix indicate stronger relationships. Loewner ordering of the individual blocks likewise indicates that dangling transcript assignment captures more variability post-Sparcle. This additional variability can be exploited to refine cell type assignment and improve gene-level analyses.

### Sparcle generalizes to additional imaging modalities

To test the generalizability of Sparcle, we examined public single molecule FISH (smFISH) images of the adult mouse primary visual cortex, comprising 3500 cells and 1,074,000 mRNA spots from 22 unique genes (15) from which we subsampled 250,000 mRNAs (154,000 within cells (61.8%) and 95,000 dangling (38.2%)). For comparison purposes, we used matching scRNA-seq data comprising 43,498 cells and 727 highly variable genes, including the 22 smFISH genes. We clustered count matrices and annotated smFISH cells and clusters (both before and after Sparcle) based on the four cell types identified by scRNA-seq (**Fig. 4a** and **Fig. S3**). Sparcle recovers 32.2% of the total mRNA across three iterations (26.3% in iteration 1, 4.7% in iteration 2 and 1.2% in iteration 3) (**Fig. S4**). Analysis of Gramian matrices reveals that post-Sparcle cell-type covariances are more similar to the matching scRNA-seq cluster covariances (Box’s M test, p > 0.05; Frobenius norm closer to 0) (**Fig. 4b**) and the cluster-cluster cross covariance matrix between scRNA-seq and MERFISH has a stronger diagonal matrix post-Sparcle (**Fig. 4c**). Cluster analysis shows that Sparcle assigns multiple genes to at least one additional biologically relevant cell type; for example, *Kcnk2, Sv2C* and *Thsd7a* belong to either endothelial or excitatory cells pre-Sparcle, and expand to both cell types after Sparcle (**Fig. 4d**).

**Figure 4.**
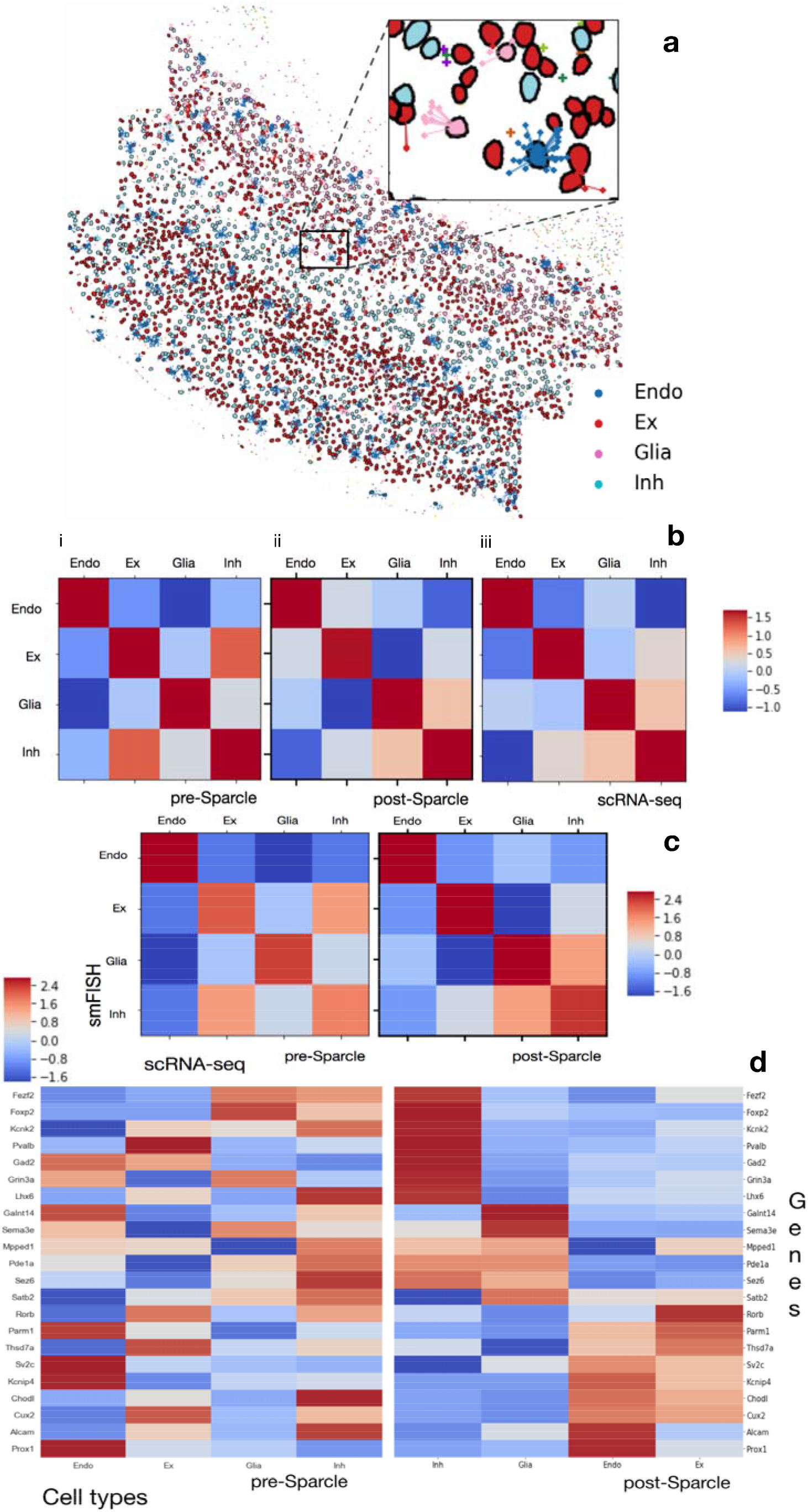
Sparcle generalizes to smFISH data. **a.** Single smFISH FOV of mouse visual cortex containing 3500 cells and 250,000 randomly subsampled mRNAs; of 93,000 dangling transcripts, 3300 are shown for visual clarity. Cells and connections to dangling mRNAs from 22 genes are colored based on matching scRNA-seq clusters. Inset, magnification. **b.** Cluster-cluster covariance (Gramian) matrices between (i) smFISH, (ii) smFISH with Sparcle and (iii) scRNA-seq indicate that Sparcle-treated count matrices are more similar to scRNA-seq (Box’s M test for pre-Sparcle vs. scRNA-seq median χ2 (2, N = 3500) = 0.36, p = 0.48, Frobenius norm = 9.42; Box’s M test for post-Sparcle vs. scRNA-seq median χ2 (2, N = 3500) = 0.92, p = 0.78, Frobenius norm = 1.83). **c.** Cluster-cluster cross covariance matrices between scRNA-seq and MERFISH clusters pre- and post-Sparcle indicate stronger within-cluster variances (dominant diagonal) after Sparcle. **d.** Expression of 22 smFISH genes (rows) within each cluster (columns) pre- and post-Sparcle, arranged by hierarchical clustering of the post-Sparcle matrix. Sparcle generates clearer blocks of gene expression per cluster and improves assignments for all four cell types. Color scale in all panels represents z-scored values of log-normalized expression per cluster.

## Discussion

Sparcle addresses an important need in the rapidly evolving field of single-cell spatial transcriptomic imaging—the recovery of transcripts outside of segmentation cell boundaries. Our approach assigns dangling transcripts to cells based on their gene expression similarity, through an imaging-platform-independent and comprehensive probabilistic model.

Sparcle takes a conservative approach to dangling mRNA assignment since there is no ground truth for cell segmentation, and many cell morphologies are very complex; for example, neuronal cell bodies are typically partially segmented, whereas mRNA-bearing axons are not usually segmented at all. On MERFISH and smFISH images of brain sections, Sparcle recovers more than 50% of dangling mRNAs, driving improved cell type calls and gene-level analyses. We have shown that Sparcle mRNA assignments lead to refined count matrices and improved assessment of cell type heterogeneity.

Spot-based Spatial cell-type Analysis by Multidimensional mRNA density estimation (SSAM) (16) and Bayesian Segmentation of Spatial Transcriptomics Data (Baysor) (17) are recent quantification approaches that aim to circumvent the need for cell segmentation. Like Sparcle, Baysor works by assigning transcripts to artificially generated mock cells. However, Baysor requires that a minimum number of transcripts per cell be specified a priori. The approach assumes that cells consist of homogenous transcripts (whereas cells and their neighbors can exhibit a diverse mix of mRNAs, e.g. **Fig. 1c**), that they have a fixed number of mRNA neighbors (whereas Sparcle allows neighborhood counts to vary within a fixed neighborhood size), and that cells take on an ellipsoid shape through a bivariate normal prior (not the case for many neuronal cells, for example).

Sparcle relies on imperfect region-independent cell segmentations as a starting point, and it uses MLE to assign transcripts to inferred cell clusters. Given the high number of dangling mRNAs, this a relatively simple, computationally cheap and powerful strategy that does not require customization for different imaging protocols. In contrast, Baysor requires that its core EM machinery be tailored to each imaging protocol.

A crucial assumption for SSAM is the uniform distribution of mRNA within a cell, which is unrealistic given the diversity of mRNA within a cell (**Fig. 1c** and **Fig. S1**). SSAM further requires the approximation of an average cell diameter, which must be adjusted for every imaging scenario. This also means that the width of the Gaussian kernel used in the KDE needs to be chosen based on the imaging protocol. A wrong choice of bandwidth can lead to signal contamination from adjacent cells.

Transcript assignment by Sparcle can be further improved. Sparcle currently utilizes an MLE protocol which does not assume an informative prior distribution for the dangling mRNAs. For any given gene, we can introduce a prior distribution for the localization of its transcripts and implement a maximum a posteriori (MAP) Bayesian inference procedure based on their spatial diversity. Additional improvements can be achieved by staining for cellular membranes in addition to nuclei, to provide rough boundaries for entire cells. This improved segmentation would help to resolve dangling transcripts positioned farther from the nucleus. A software version in a language such as Java may also run more efficiently.

Sparcle is shown to effectively assign dangling mRNAs to cells and correct the count matrices. We hope that the computational infrastructure provided by Sparcle will further contribute to the emerging landscape of high-throughput spatial transcriptomic image analysis.

## Methods

### Algorithmic overview

As input, Sparcle takes approximate cell segmentations generated using any method for an FoV. The cell segments across all the FoVs are collected, and Sparcle creates a global count matrix of cells and genes, which is clustered to identify cell types. The large number of transcripts in each cell justifies modelling gene expression as a Gaussian distribution, based on the central limit theorem assumption. An image can thus be considered as a mixture of cell-specific Gaussian distributions, which can be used in a Dirichlet process mixture model (DPMM) for identifying heterogeneous cell types. First, Sparcle learns the cluster-specific first and second order moments (mean and covariance) of each cluster. Next, for each dangling mRNA, a mock cell is created using mRNA transcripts occurring within a certain radius of the dangling mRNA. The mock cell is compared to previously computed cluster-specific moments with the objective of maximizing the likelihood of a dangling mRNA being similar to a cluster and then assigning the mRNA to the closest neighboring cell belonging to the most similar cluster. Neighboring cells are calculated based on Euclidean distances between the mRNA and cell centroids. As each mRNA gets assigned to a cell, we update the count matrix. At the onset of each subsequent iteration, the updated global count matrix is re-clustered, the cluster-specific moments are recomputed and the assignment of dangling mRNAs continues. Through this construction, we are, in essence, building a global model with global cluster-specific means and covariances that all the FoVs have contributed to and will therefore adhere to. Using this global approach, we can ensure that the underlying probabilistic model is constant across FoVs per iteration as well as across iterations.

If we cluster each FoV separately in order to build a local structure, it becomes necessary to ‘match’ the clusters between FoVs. This can be both computationally intensive and unstable, because not all cell types are present in all FoVs, and thus, clusters can differ across FoVs. To build the global structure, we need to perform roughly 400 x k x k’ pairwise stable matching of clusters between FoVs, where k and k’ are the number of clusters for any given pair of FoVs. These comparisons would be riddled by missing or unseen clusters, leading to an improper global structure.

Sparcle therefore resorts to constructing a global structure of cell types from the beginning, to ensure a reliable set of clusters to which dangling mRNAs can be assigned. Sparcle comprises several canonical building blocks, including clustering to identify cluster-specific moments, constructing a mock cell for each dangling mRNA to enable comparison of a dangling mRNA to cluster moments, and maximum likelihood estimation.

### Mockcell window size and shape choice

The mockcell is designed as a circle centered at the dangling mRNA. We consider a circular window to approximate the cell body of an average neuronal cell. To estimate the approximate radii of two mockcells A and B, we draw bounding boxes around those cells (see **Fig. S2** for example corresponding to calculations below). The area of a cell’s bounding box = (*x_1_* – *x_2_*) * (*y_1_* – *y_2_*). We equate this to the area of a circle = *πr^2^*, to estimate *r*.

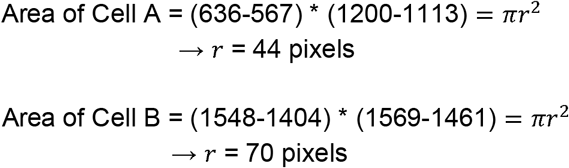

Since most cells in MERFISH and smFISH data have *r* <= 75 pixels, we have set *r* = 80 pixels as the default radius for a mockcell.

### Weighted mockcell creation

Every dangling mRNA is represented as a mockcell and is a vector of length equal to the number of genes, G. Using the circular window of radius *r* centered at the dangling mRNA, a mockcell is constructed using an inverse distance weighting (IDW) method where distances from the dangling mRNA (or the center) to each of the neighboring mRNA falling within *r* are interpolated to create a weighted average for each g in G. With this construction, mRNAs closest to the dangling mRNA have more influence than those farther away. The weighted average *u_g_* for each gene *g* for the mockcell centered at *x* is calculated based on the simplest weighting function or Shepard’s method (18) that uses the weights in inverse power as:

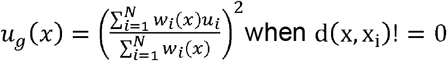

and
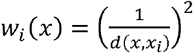 where 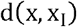 is the distance between the interest point *x* and a point in the neighborhood *x_i_* (denoting a neighboring mRNA) where there are *i* = 1, …*N* observations of gene *g* in the neighborhood and *w_i_*(*x*) is the weight or predicted value at point *x*.

We use a modification of Shepard’s method that calculates the interpolated value using only nearest neighbors within the r-sphere (instead of the entire sample space). Weights in the above equation are modified in this case to give 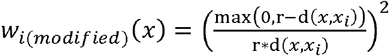 that are used to compute the modified weighted average *u_g_mod_*(*x*) which is entered for each *g* the mRNA accounts for in the mockcell.

### Spatial distances and how they are incorporated into Sparcle

By orienting every image to a 2D Cartesian coordinate system, we assume each image to reside on a Euclidean plane. This allows us to calculate the inter-cellular distances using pairwise Euclidean distances between cell centroids. Further, the local concentration of mRNA transcripts within a mockcell is derived by computing the Euclidean distances between the transcripts and the mockcell’s centroid.

### Maximum likelihood method

We assume that the gene expression in each cell follows a multivariate Gaussian distribution. This enables us to view an image as a mixture of cell-specific Gaussian distributions, and to use a Dirichlet process mixture model (DPMM) for clustering the cells into *K* different cell types.

In order to assign a dangling mRNA, *m*, to one of the inferred cell types *k* = 1 … *K*, we use the maximum likelihood (ML) method. Assuming there are *k* cell types, the mockcell originating at a dangling mRNA, is tested for each cell type’s parameters i.e. multivariate mean *μ_k_* and covariance ∑*_k_* to identify *k_ML_* that maximizes the likelihood distribution. We assume the likelihood distribution to be that of a multivariate Gaussian. The analytic form of the ML method can be written as:

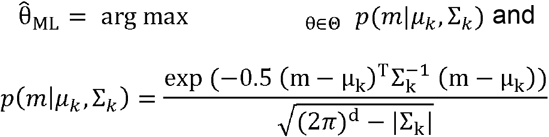

where 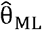 is the maximum likelihood estimate of the Gaussian parameters *θ* = [*μ_k_,∑_k_*], Θ is the finite-dimensional parameter space for *θ, d* is the number of genes and *p*(*m*|*θ*) is the non-degenerate case of the multivariate Gaussian probability density function (pdf). We ensure the non-degenerate pdf by constructing positive (semi) definite ∑*_k_* using Givens rotations as described in the next section. The dangling mRNA is assigned to the nearest neighboring cell belonging to the closest ‘likely’ cell type, *k_ML_* described by 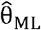. The count matrix entry for the cell identified and gene associated by the dangling mRNA is updated. The assigned dangling mRNA is removed from the candidate list of mRNAs to be assigned. The ML method is known to be optimal for large data sizes (19).

### Givens rotations on the covariance matrices

To ensure that covariance matrices used in the maximum likelihood density estimation are positive semi-definite, and to avoid unnecessary cost overhead by inverting these matrices, we perform QR decomposition using the Givens rotations on the covariance matrices. These are efficient transformations on sparse matrices and can be better parallelized as well.

For any real square matrix *A*, the QR decomposition is *A = QR* where *Q* is an orthogonal matrix and *R* is an upper triangular matrix. QR decompositions are computed with a series of Givens rotations. In each rotation an element in the subdiagonal of the matrix is zeroed out, to form the symmetric R matrix. The product of all the Givens rotations forms the orthogonal Q matrix.

In practice, Givens rotations are not actually carried out by building the entire matrix to perform the matrix multiplication between *Q* and *R*. Instead, only the upper triangle of *R* is built out and multiplied with *Q* during each Givens rotation which is equivalent to a sparse Givens matrix multiplication, without having to explicitly handle the sparse elements. This Givens rotation procedure is useful when only a relatively few off diagonal elements need to be zeroed out implying the matrix is still dense, and can be more easily parallelized (20, 21).

### Runtime complexity analysis

We calculate the Big O notation for an entire run of Sparcle for one MERFISH FOV. On average, a MERFISH FOV consists of 80 cells, of which ~50% of mRNA transcripts fall within a cell and the rest are dangling mRNA. We consider n the number of cells, *d* the number of genes, *m* the number of dangling mRNA transcripts in the first iteration, *m’* the number of dangling mRNA transcripts in the subsequent iterations, x the number of line segments constituting each cell boundary, *k* the number of mRNA neighbors per dangling mRNA for constructing the mockcell and *c* the total number of iterations. We have observed that setting *c* to 3 is usually sufficient since most of the dangling mRNAs are assigned in iterations 1 and 2.

1. Construct the count matrix for each FOV and consolidate to an overall count matrix: *O*(*n*), 1 min
2. Iteration1: O(*nd*^2^) + nO(n) + mO(k) = O(*nd^2^*), 3 mins

a. Maximum likelihood estimation calculations: *O*(*nd^2^*)
b. Euclidean distance to find neighboring cells for cell *i*: O(n)

i. n subtractions for (cell_centroid_i - neigh_cell_centroid)
ii. n squares of (i)
iii. n-1 further additions to add (ii)
iv. Final one square root. (So each of these is (at most) linear in n, and hence so is the whole algorithm.)
c. Perform *b* for n cells = nO(n)
d. Euclidean distance to create j^th^ mock cell using neighboring mRNA for j^th^ dangling mRNA: O(k)

i. *k* subtractions for (dangling mRNA_centroid_j - neigh_mRNA)
ii. *k* squares of (i)
iii. *k-1* further additions to add (ii)
iv. Final one square root. (So each of these is (at most) linear in *k*, and hence so is the whole algorithm.)
e. Perform *d* for *m* cells = *mO*(*k)*
3. Iteration *c*: O(*nd^2^)*, 2.5 mins
4. Final Iteration (combining all the assignments and plotting): *O*(*nm*) + *cO*(*m*), 1.5 mins

Therefore, overall runtime complexity is O(n) + O(*nd^2^*) + cO(*nd^2^*) + O(*nm*) + cO(*m*) ~ O(*nd^2^*) + O(*nm*) ~ *O*(*nm*) ~ *O*(*m*)

The runtime is given based on Sparcle runs on an HPC.

### Evaluating cell populations identified by Sparcle using matching scRNA-seq

a. The Gramian matrix across clusters for the pre-Sparcle count matrix, post-Sparcle count matrix and matching single-cell RNA-seq data is shown in **Fig. 2b**. For this, we compute the Box’s M test which is a parametric test to check the homogeneity of variance (homoscedasticity). Specifically, it tests for the homogeneity of covariance matrices describing multivariate Gaussian data according to one or more groups. The groups in our setting are pre-Sparcle, post-Sparcle and scRNA-seq. The test compares the product of the log determinants of each of the covariance matrices to the log determinant of the pooled covariance matrix, similar to a likelihood ratio test (22). The generated test statistic is called Box’s M statistic and is approximated using a chi-square goodness of fit. The null hypothesis for this test is that the observed covariance matrices are equal across groups. This means a non-significant test result (i.e. one with a large p-value) indicates that the covariance matrices are not different (23, 24). The Box’s M test between

- Pre-Sparcle and scRNA-seq: median chi-squared = 0.2556, df = 2, p-value = 0.3091.
- Post-Sparcle and scRNA-seq: median chi-squared = 0.9458, df = 2, p-value = 0.6201.

> A p-value greater than 0.05 indicates the variances are homogeneous and higher p-values indicate more homogeneous variances depicting that post-Sparcle and scRNA-seq covariances are more similar. Further, we check the distance between covariances using the Frobenius norm (discussed in the next section).

a. The Cluster-cluster cross covariance matrix between pre-Sparcle clusters with scRNA-seq clusters and between post-Sparcle clusters with scRNA-seq clusters (Right panel) is shown in **Fig. 2c**. We observe that in the post-Sparcle covariance,

a. the diagonal shows stronger variances **within** clusters than the pre-Sparcle covariance matrix, especially for nonneuronal cell types.
b. the off-diagonal elements indicate *weaker to no* covariance **between** clusters as opposed to the pre-Sparcle and scRNA-seq covariance. This is a preferred covariance pattern that signals stronger relationships within and weaker relationships between cell types. This means that cell types have garnered relevant mRNA transcripts via Sparcle assignments.
b. We map canonical genes per cluster for both pre-Sparcle and post-Sparcle cell types in **Fig. 2d**. We see clear and improved cluster assignments for most of the post-Sparcle clusters.

### Distance between covariance matrices using the Frobenius norm

Consider a m × m covariance matrix *A* and the set *B* with all m × m positive definite covariance matrices having similar structure (for example, uniform covariance structure). In order to find the discrepancy between *A* and the set *B*, we define 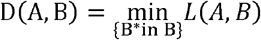 where *L(A,B)* is a measure of the distance between the two m×m matrices *A* and *B*. We consider *A* the covariance matrix from the scRNA-seq measurements and the set *B* to consist of the pre- and post-Sparcle covariance matrices that originate from the same measurement space. The matrix *B\** which has the smallest discrepancy can be viewed as that with the most likely or closest to the underlying structure of *A*. We consider the distance between the two matrices to be the square of the Frobenius-norm, or the F-norm: *L*(*A,B*) = *tr*(*(A – B*)*^T^* (*A – B*)), which is defined as the square of the sum of the absolute squares of its elements (25).

For **Fig. 2b**, the Frobenius norm for the Gramian matrices between

- Pre-Sparcle and scRNA-seq is 15.18
- Post-Sparcle and scRNA-seq is 2.34 showing that the post-Sparcle covariance is closer to the scRNA-seq covariance.

### Loewner ordering of matrices

One way to compare matrices is by using the positive definiteness (pd) or positive-semi definiteness (psd) property. We extract the block matrices from the Pearson’s correlation matrix (**Fig. 3**) and test them for pd or psd using the Cholesky decomposition. Cholesky decomposition factors a psd matrix *A* into: *A* = *LL^T^* where *L* is the lower triangular matrix and also the Cholesky factor of *A*. If L is invertible, then A is a pd matrix. Since the blocks are from covariance matrices, the blocks satisfy either the pd or psd property. Assume there are two covariance matrices (or multivariate variances) *C* and *D* where *C* and *D* are by definition pd or psd or verified using the Cholesky decomposition described above. If upon removing *D* from *C* (i.e. *C – D*) we get a positive (semi) definite matrix that means that *C* has captured *more* viable variability within the system. We use this Loewner ordering of covariance matrices to show that the post-Sparcle covariance matrices capture additional real variability which went unaccounted for in the pre-Sparcle covariance matrices.

## Declarations

### Ethics approval and consent to participate

Not applicable

### Consent for publication

Not applicable

### Availability of data and materials

MERFISH data and corresponding scRNA-seq data can be found at (9). For MERFISH, there are 400 FOVs, each ~3GB and 4.1 Megapixels in size. There are 31,241 cells across the 400 images of which 2.7 million mRNA transcripts fall within cells and 2.8 million are dangling mRNA. The matching scRNA-seq data has 31,299 cells and 27,998 genes.

For smFISH, we utilise a single image that was stitched and provided by Brian Long from the Allen Brain Institute. The image is that of the primary visual cortex (VISp) region of an adult mouse brain. There are 3500 cells and 22 genes. There were 1074,000 mRNA within this image of which we subsample 250,000 mRNA of which 154,000 mRNA are within cells and 95,000 are dangling mRNA that is used by Sparcle. The matching scRNA-seq data had 43498 cells and 45,000 genes and is available at (15).

The open-source, platform independent software implementation of Sparcle is available on Github: https://github.com/sandhya212/Sparcle_for_spot_reassignments

Code is written in Python 3 and is parallelized to process multiple FOVs during each iteration. We have also developed code that is necessary to reconstruct each FOV from the stage coordinates to image coordinates, plot the cell segment boundaries obtained from the segmentation, and plot the mRNA within cells and dangling mRNAs.

### Author’s contributions

SP, DP conceived the study and designed the concept and idea of SPARCLE. SP developed the code for SPARCLE. SP acquired and performed data analyses on MERFISH and smFISH data. TN provided critical comments and discussions. DP supervised the study. All authors wrote and approved the final manuscript.

## Acknowledgements

We thank Jeffrey Moffitt (Harvard University) for sharing MERFISH data and for in-depth discussions on biological interpretation of the data. We also thank Brian Long and Jeremy Miller for sharing the Allen smFISH data and thoughts on the matching single-cell RNA-seq data.

## Supplementary Figures

**Figure S1:**
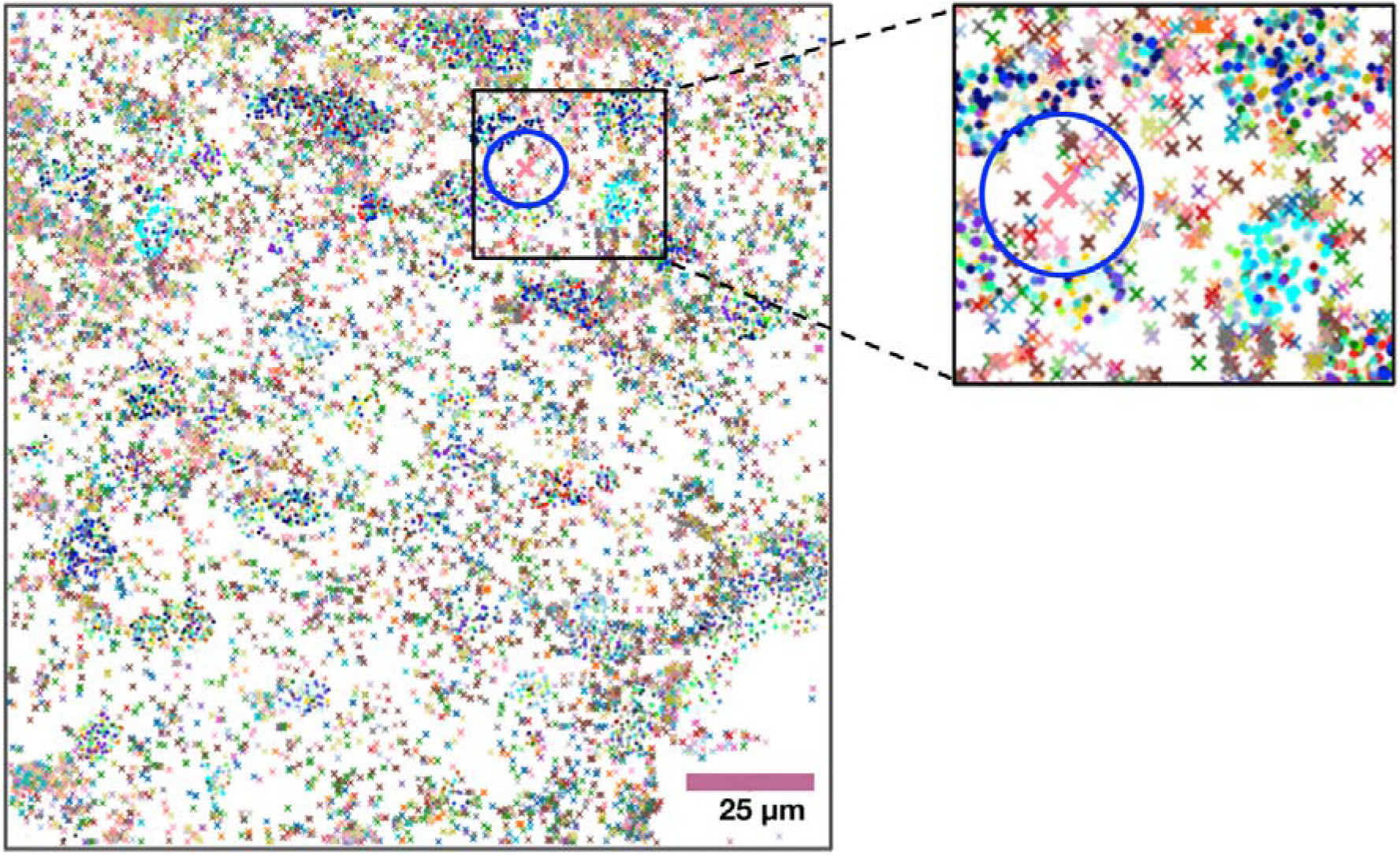
Weighted Mockcell creation. A MERFISH FOV showing spatial allocation of 134 genes in a section of the mouse hypothalamus optic region [9]. Dots represent mRNA within cell segmentations, colored by cell cluster assignment. Crosses represent unassigned (dangling) mRNA. Inset, a zoomed-in region showing mock cell (blue circle), centred at a dangling mRNA (pink cross). All mRNA (dots and crosses) within the circle will contribute to that mock cell, weighted inversely to the distance from the central mRNA.

**Figure S2:**
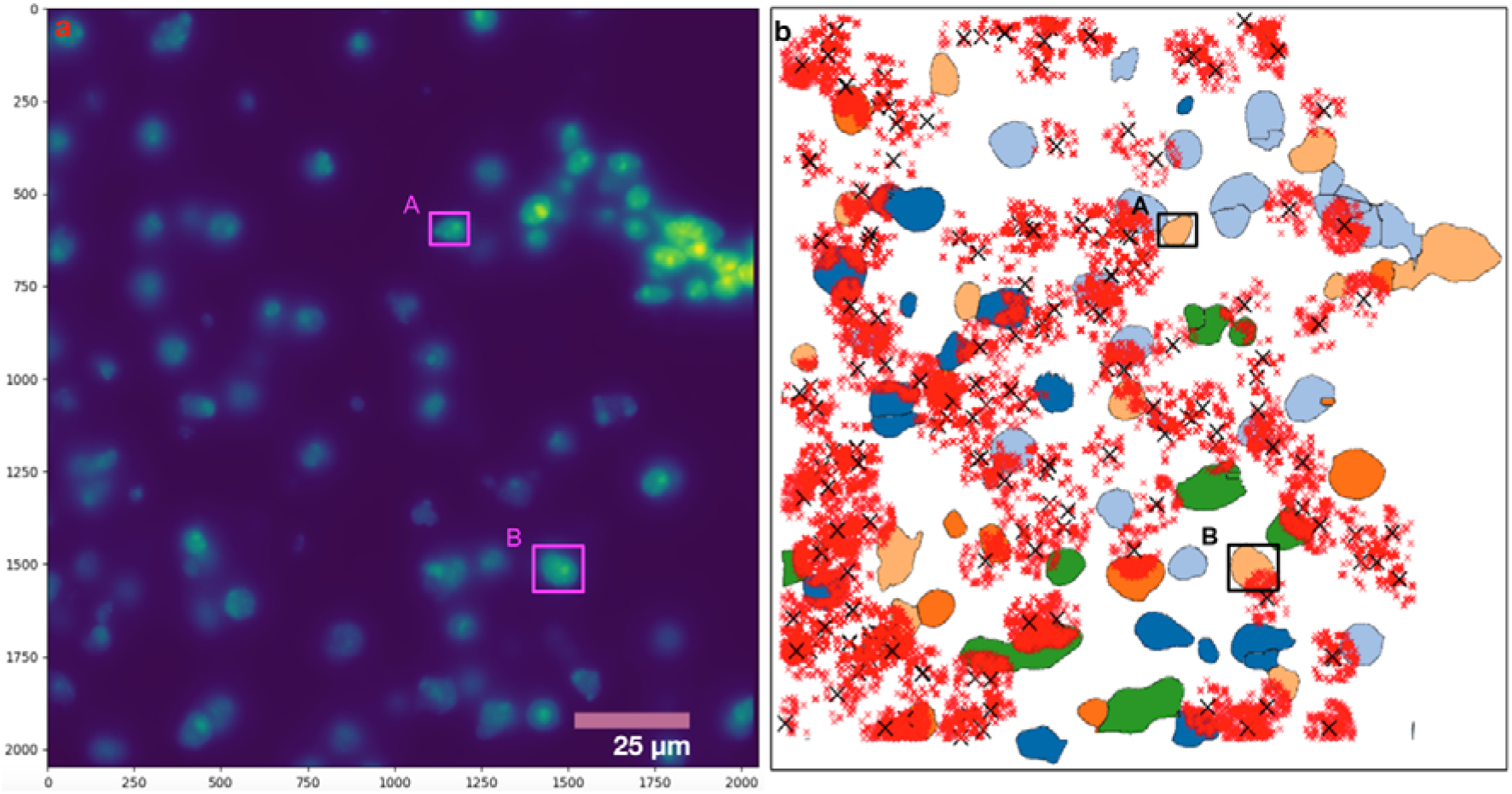
Mockcell creation. **a**. A MERFISH image shown in image coordinates after perspective projection from stage coordinates. This projection allows the x and y axis to start from (0,0) and enables the calculation of Euclidean distances between cell centroids and mRNA transcripts. We use this to estimate the radius r for a mockcell. For example, we draw a bounding box around cell A and cell B and observe that its radius is around 80 pixels. This is used as a proxy for the mockcell creation. **b**. The same MERFISH image recreated in Python using cell segmentation coordinates to outline the neuronal cells. The cells are colored based on their cell type. A dangling mRNA is shown as a black cross with red crosses, traced within a circle of radius 80 pixels, identifying those mRNA that will contribute to its mockcell.

**Figure S3:**
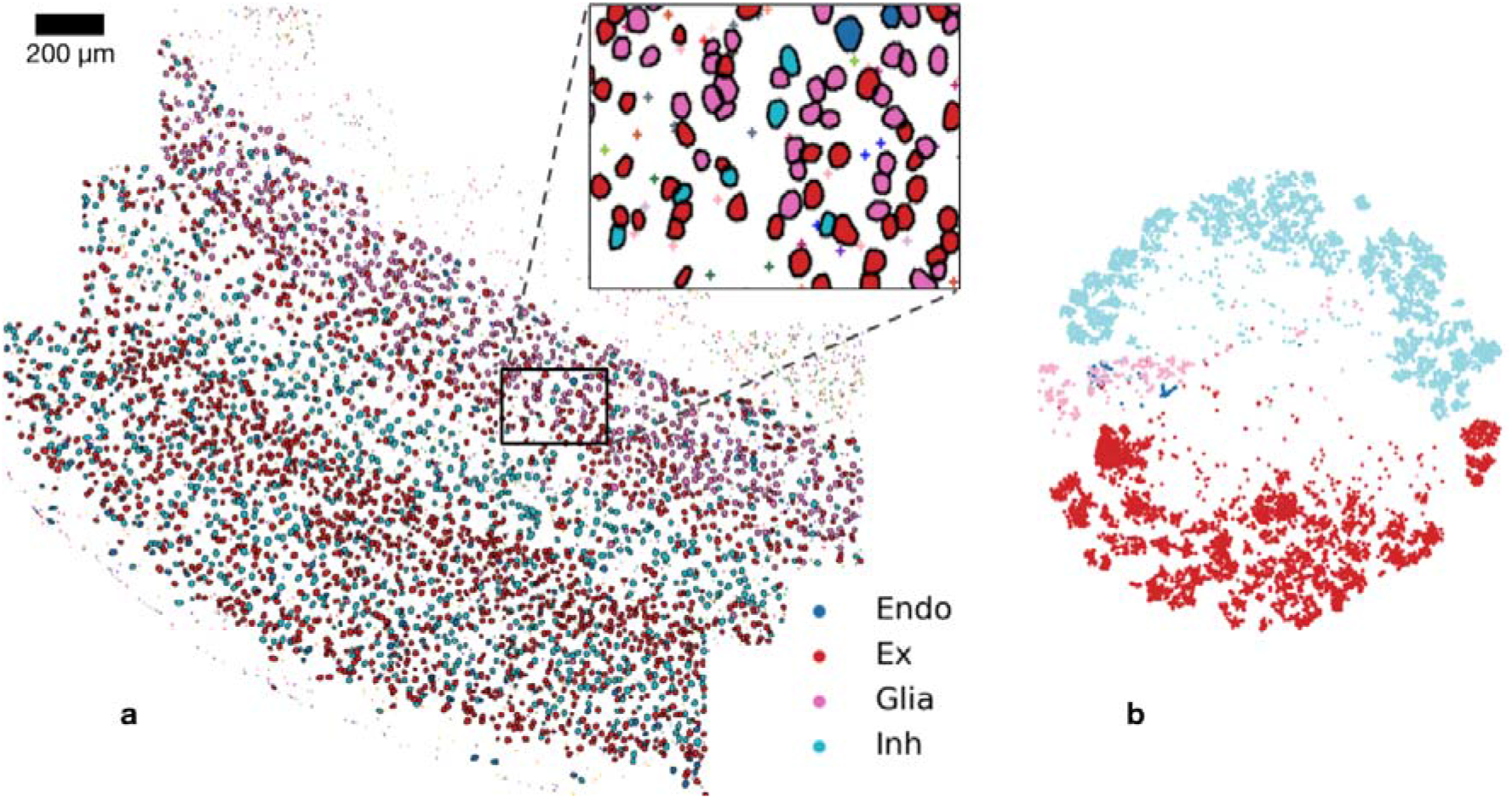
Allen smFISH data and matching scRNA-seq data. **a**. Single smFISH FOV of mouse visual cortex containing 3500 cells and 250,000 randomly subsampled mRNAs; of 93,000 dangling transcripts, 3300 are shown for visual clarity. Cells and dangling mRNAs are colored based on matching scRNA-seq clusters. Inset, magnification with dangling mRNAs (+ symbols). The dangling mRNAs account for 22 smFISH genes: *Alcam, Chodl, Cux2, Fezf2, Foxp2, Gad2, Galnt14, Grin3a, Kcnip4, Kcnk2, Lhx6, Mpped1, Parm1, Pde1a, Prox1, Pvalb, Rorb, Satb2, Sema3e, Sez6, Sv2c, Thsd7a*. **b**. tSNE projection showing proportions of the four hierarchical clusters corresponding to endothelial (Endo), excitatory (Ex), glial (Glia) and inhibitory (Inh) cell types of the matching scRNA-seq data, which comprises 43,498 cells and 45,000 genes.

**Figure S4:**
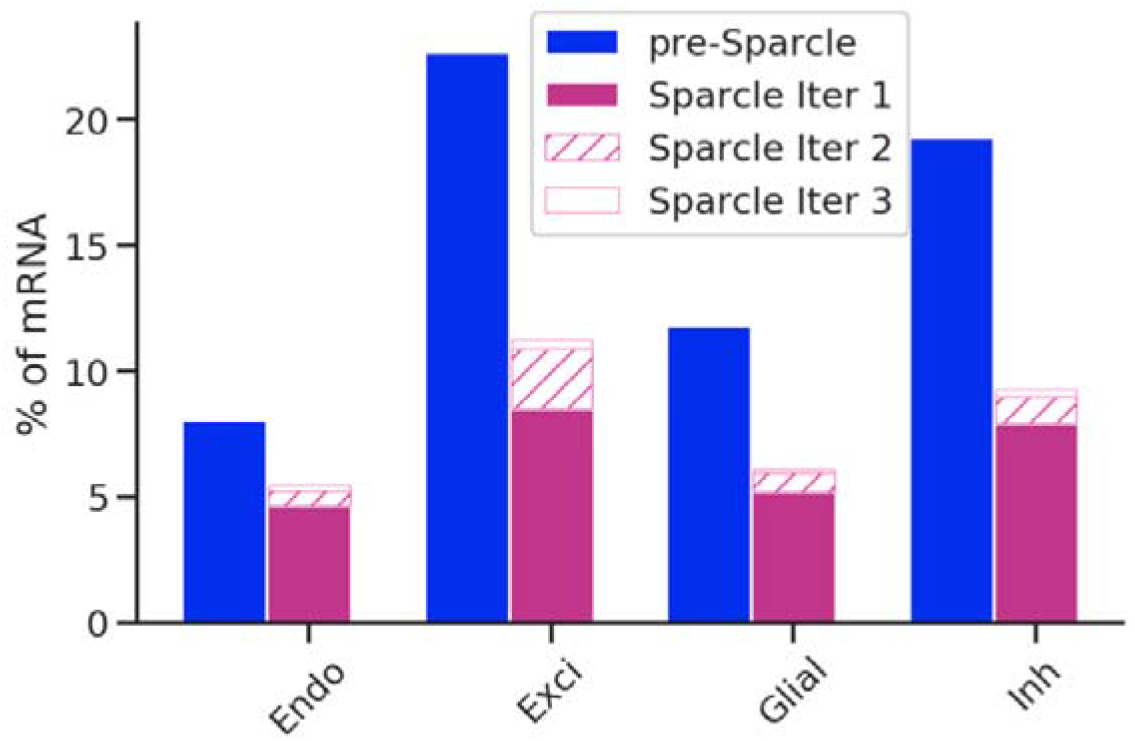
Percentage of mRNA recovery across the 4 dominant cell types using Sparcle in the Allen smFISH data. We subsample 250,000 mRNAs from a total of 1,074,000 within this image; 154,000 mRNAs (61.8%) are within cells and 95,000 (38.2%) are dangling. Cell type proportions of within-cell mRNA in the smFISH data are in blue (endothelial 8.1%, excitatory 22.7%, glial 11.8%, inhibitory 19.3%), and additional dangling transcripts assigned by Sparcle are in pink, comprising 32.2% of the total mRNA (endothelial 5.5%, excitatory 11.25%, glial 6.1%, inhibitory 9.3%) across three iterations; only 6% of the total mRNA remains unassigned.

